# Learning in the eyes: specific changes in gaze patterns track explicit and implicit visual learning

**DOI:** 10.1101/2020.08.03.234039

**Authors:** József Arató, Constantin A. Rothkopf, József Fiser

**Author notes:** Current Affiliation: Cognitive Science Hub, University of Vienna, Liebiggasse 5, Vienna 1010, Austria (József Arató). Correspondence: József Fiser, Department of Cognitive Science, Center for Cognitive Computation, Central European University, Nádor utca 9, 1051 Budapest, Hungary.

## Abstract

What is the link between eye movements and sensory learning? Although some theories have argued for a permanent and automatic interaction between what we know and where we look, which continuously modulates human information- gathering behavior during both implicit and explicit learning, there exist surprisingly little evidence supporting such an ongoing interaction. We used a pure form of implicit learning called visual statistical learning and manipulated the explicitness of the task to explore how learning and eye movements interact. During both implicit exploration and explicit visual learning of unknown composite visual scenes, eye movement patterns systematically changed in accordance with the underlying statistical structure of the scenes. Moreover, the degree of change was directly correlated with the amount of knowledge the observers acquired. Our results provide the first evidence for an ongoing and specific interaction between hitherto accumulated knowledge and eye movements during both implicit and explicit learning.

## Introduction

Across their lives, people make 2-3 saccades per second during their wake period, which fundamentally determines the sensory information reaching their conscious cognition. Yet, despite an extended literature on the control of eye movements(Findlay & Gilchrist, 2003; Hayhoe & Ballard, 2005; Kowler, 2011; Yarbus, 1967), we have only a rudimentary understanding of how past experiences influence the deployment of attention as indexed by eye movements(Wolfe & Horowitz, 2017). These include observations that gaze biases can emerge from a lifetime of experience, such as taking the inherent uncertainty of the visual system into consideration during visual search(Najemnik & Geisler, 2005), anticipating a ball’s trajectory in sports(Brockmole & Henderson, 2006; Land & McLeod, 2000), the tendency to perform visual search from left to right(Spalek & Hammad, 2005), using learnt semantic knowledge(Võ & Wolfe, 2013) or meaning in real world scenes (Henderson et al., 2018). At shorter time-scales, object co-occurrences(Brockmole & Henderson, 2006; Mack & Eckstein, 2011) and episodic memory have been shown to guide visual search(Li et al., 2018). Past experience on an even shorter time-scale can also influence gaze selection, for example when integrating visual information in a given scene with what has been learned about stimulus statistics within minutes(Hoppe & Rothkopf, 2016; Yang et al., 2017). A number of these studies investigate jointly how humans develop specific eye movement patterns based on experience with the structure of sensory input and how they use specific eye movement strategies to solve particular tasks(Brockmole & Henderson, 2006; Hoppe & Rothkopf, 2016; Land & McLeod, 2000; Li et al., 2018; Mack & Eckstein, 2011; Nelson & Cottrell, 2007; Yang et al., 2017). However, all the above studies considered specific tasks (e.g. categorization, search), and they focused on end results, that is, they showed that after practice, observers learned the identity and/or location of diagnostic features of the task, and their eye movements became more related to these features. Such studies do not clarify, which of the two competing alternatives best describes the nature of the interaction between acquired knowledge and eye movements in everyday life. First, this interaction could emerge only within the specific context of a clearly defined task via top-down control on sensory processing by high-level explicit knowledge established earlier. Alternatively, the interaction could be a general and ongoing process that modulates human information-gathering behavior all the time and continuously supports both implicit and explicit learning. These two alternatives have very different consequences on how dynamic active perception and the role of eye movements within such perception should be framed with major implications on the relationship between perception and cognition

In this study, we address two questions that help evaluate these two alternatives. First, we asked whether there is a difference between how eye movements and sensory learning interact during an explicit task vs. in a task-free observation of structured sensory stimuli. Second, we assessed whether the effect of learning on eye movements is manifested immediately and proportionally with the amount of learned knowledge regardless of this knowledge being implicit or explicit, or alternatively, the effect emerges only after the acquired knowledge becomes explicitly accessible. To explore these issues, we adapted the paradigm of spatial statistical learning(Fiser & Aslin, 2001), which allows investigating the process of learning under various levels of implicitness. We altered the paradigm in a gaze-contingent manner, where in each trial, observers saw only a small part of the composite display around their fixation point at a time, and thus through each fixation, they could access different segments of the underlying scene, which consisted of multiple abstract shapes in complex statistical relationships. We coupled this paradigm with either an explicit task (Exp. 1), in which the underlying general structure of the scenes was verbally revealed to the observer prior to the experiment, or under the typical implicit condition of visual statistical learning (Exps. 2&3), where observers had no task other than to explore the unknown scene without any further instructions. This setup allowed investigating, in a continuous manner, the entire process of learning the underlying structure of the scenes from the naive to the expert state, the changes in eye movement patterns during learning, and the effect of explicitness of the knowledge that the observers gathered and relied on.

We found that observers’ knowledge about the underlying structure of the scenes acquired across multiple presentations induced a specific and significant change in their eye movement patterns. This change reflected the particular spatial structure of the constituents making up the visual scenes, and it progressed proportionally to the amount of learning throughout the learning process. Remarkably, while there was a difference in learning speeds between the conditions, when observers had prior explicit vs. no explicit knowledge, there was no difference between the two conditions in terms of how much a given amount of learning altered the eye movement patterns. Because changes in learning were detectable earlier than changes in gaze patterns, this supports the view that acquired knowledge is integrated continuously into the observer’s internal representations without the need for an explicit learning context, and that this knowledge continuously contributes to the control of subsequent information gathering through influencing eye movements.

## Results

### Explicit learning of regularities influences eye-movements

To establish whether there is an ongoing link between the acquisition of complex environmental regularities and eye-movements during learning, we explicitly revealed the rules of the underlying statistical structure of the presented scenes (but not the identity of shapes in pairs) before Exp. 1. On the 2-IFC familiarity test, participants demonstrated significantly above chance performance (Fig 1D, M=70.56% 95%CI [64.94, 76.17] t(39) =7.09, *p*<.001, Cohen’s *d*=1.12), indicating that they, at least partially, acquired the underlying regularities of the training scenes. To investigate the effect of the learned underlying structure on eye-movements, we analyzed whether the exploratory and confirmatory gaze transitions were influenced by the pair structure during training through the slope of regression (β) fitted to the proportion of exploratory and confirmatory looks across trials. The proportion of both types of looks following the pair structures was steadily increasing over the trials (Exploratory: β=.0245, *p*<.001, Fig 2A; Confirmatory: β=.0301, *p*=.026, Fig 2B). Furthermore, both measures were predictive of the performance on the final familiarity test on average (Exploratory: *r(38)*=0.39, *p*=.013; Confirmatory *r(38)*=0.70, *p*<.001). Moreover, this predictive power of eye movement patterns on final test performance gradually emerged during the learning phase (Fig 3). To test whether beyond the overall influence, the specific content of learning could also be deciphered from the observer’s eye-movements, we used the orientation specific parameters (α_1-3_) of the the model-based statistical analysis to predict the observer’s performance with the differently oriented pairs during the familiarity test. This test showed clear evidence for a significant relationship between the α parameters of eye-movement modulation and learning performance with pairs in all three orientations (Fig 4 A-C). Summarizing the results of Exp. 1, we found that explicit learning of complex regularities can influence eye-movement patterns. Previous evidence on the number of fixations until finding a target (Najemnik & Geisler, 2005; Peterson & Kramer, 2001) and looking times (Hoppe & Rothkopf, 2016) suggested that eye-movements can utilize environmental regularities. Our findings extend these results by showing that, with an explicit task, the patterns of explorative eye-movements become sensitive to newly learned spatial stimulus regularities, and the change in eye-movements reflect the amount of learning.

**Figure 1.**
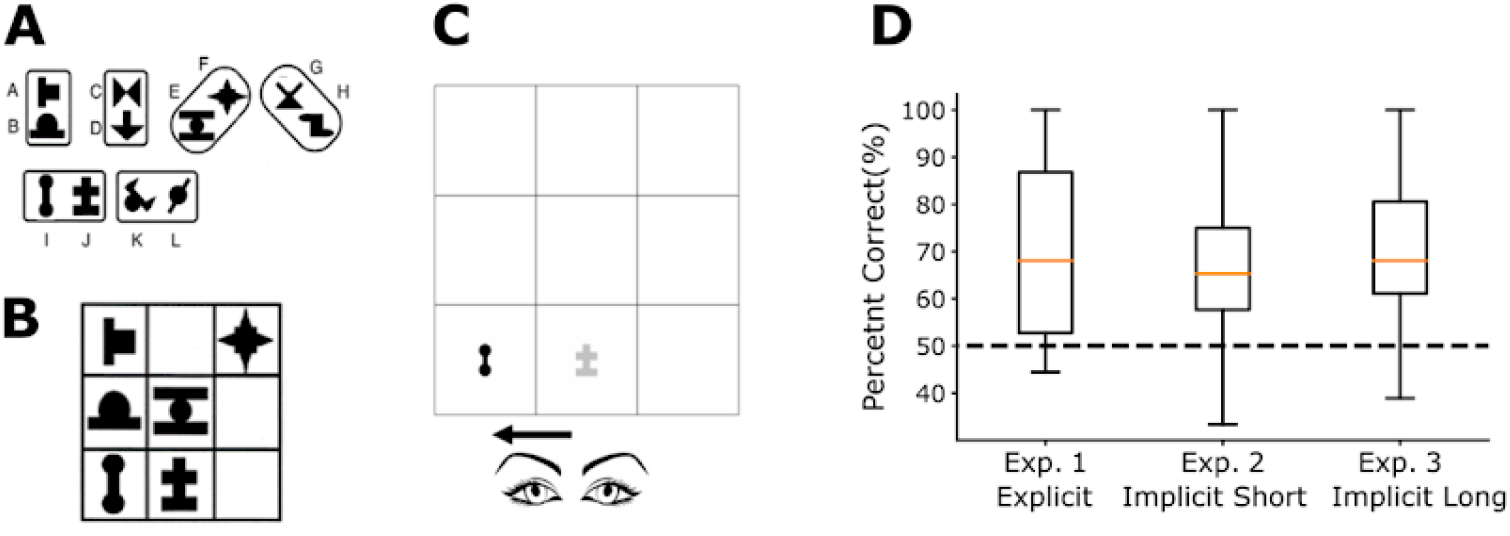
Experimental design and test results. **A)** A set of 12 abstract shapes were randomly assigned to 6 pairs (2-vertical,2-horizontal,2-diagonal) for each participant. **B)** One example of the 144 possible scenes that were assembled from 3 differently oriented pairs randomly arranged on a 3 by 3 grid following the method of previous studies of spatial statistical learning. **C)** Example trial snapshot of the gaze-contingent statistical learning paradigm applied in this paper with the underlying structure of the trial scene shown in B, while the participant’s gaze moved from the bottom middle to the bottom left cell (indicated by the arrow). **D)** Results of the 2-IFC familiarity test after the learning phase in the three experiments differing only in instructions and training lengths showed highly significant learning performance (N=40, each, Error bars: full range of data,). Test performance was not different across the three experiments (F(2,117)=0.89, p=.415, η_p_^2^=.01).

**Figure 2:**
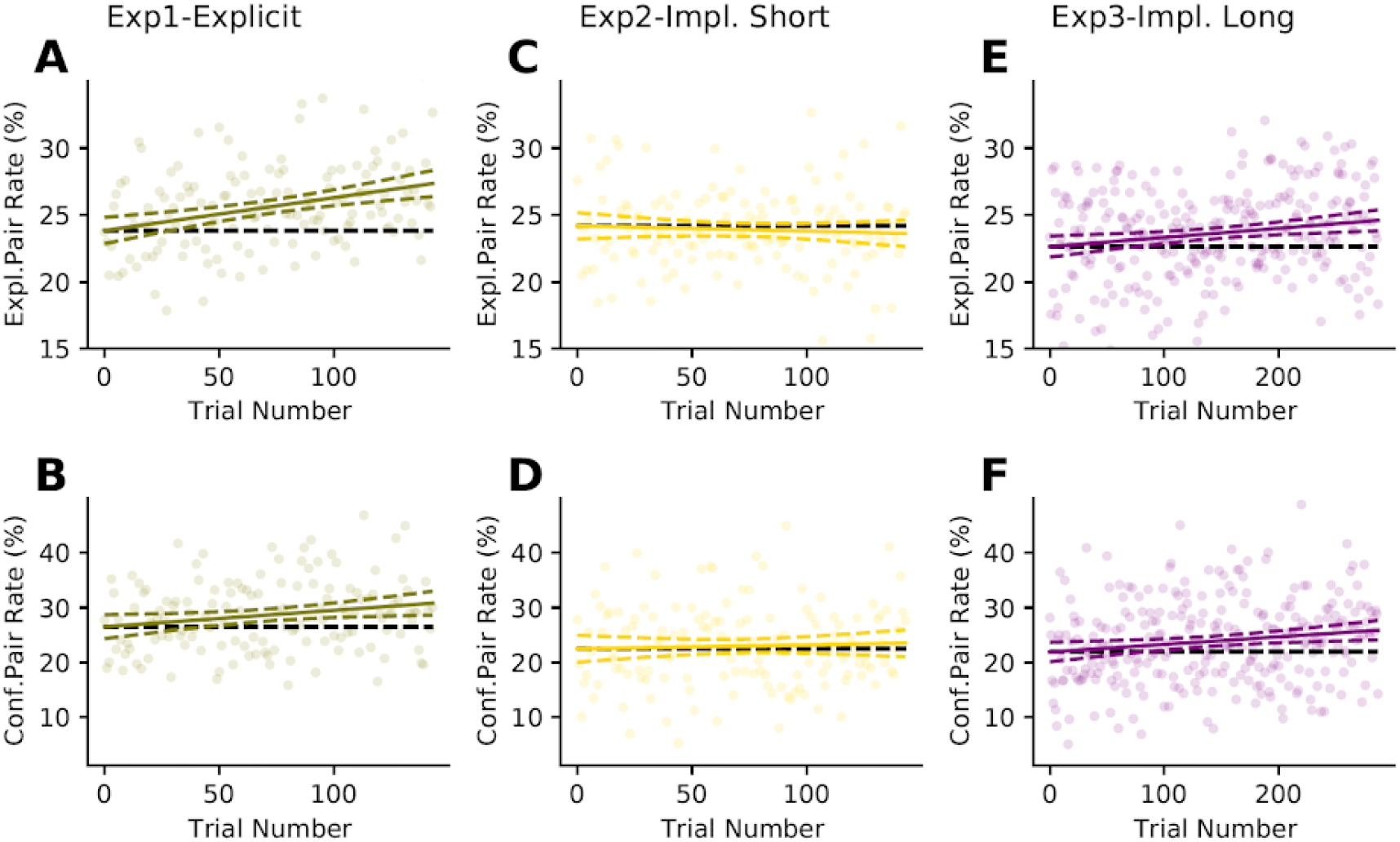
Eye-movements are progressively influenced by learned statistical regularities. Columns indicate the three experiments (Exp. 1: A,B; Exp. 2.: C,D; Exp. 3.: E,F), rows show the two measures (Exploratory and Confirmatory gaze transitions) used to quantify the relation between learned underlying spatial regularities and eye-movement patterns. Dots represent per trial proportion values for each observer for the two measurements, group performance is shown by the least squares regression line (solid) and the 95% confidence interval (dashed). Black dashed horizontal line indicates chance performance. **Top Row:** The proportion of explorative eye-movements that were performed according to the statistical structure of the scene (moving from a shape to its pair) was increasing over-time when the instructions were explicit (Exp. 1: **A**, β =0.0245, *p*<.001) or during long implicit learning (Exp. 3: **E**, β=0.0068, *p=.005*), but it stayed non-significant during the short implicit learning ( Exp. 2: **C** β =−0.0039, *p=.513)*. **Bottom Row:** The same conclusions are supported by the Confirmatory Gaze Transitions measure, the proportion of within trial returns to cells already visited on a given trial that were performed within shapes forming pairs. Again, there was a significant increase in Exp. 1 (**B**, β =0.0301, *p*=.026, solid line) and Exp. 3 (**F**, β =0.0139, *p*=.012), but no change in Exp. 2 (**D**, β =0.007, *p=.643).*

**Figure 3:**
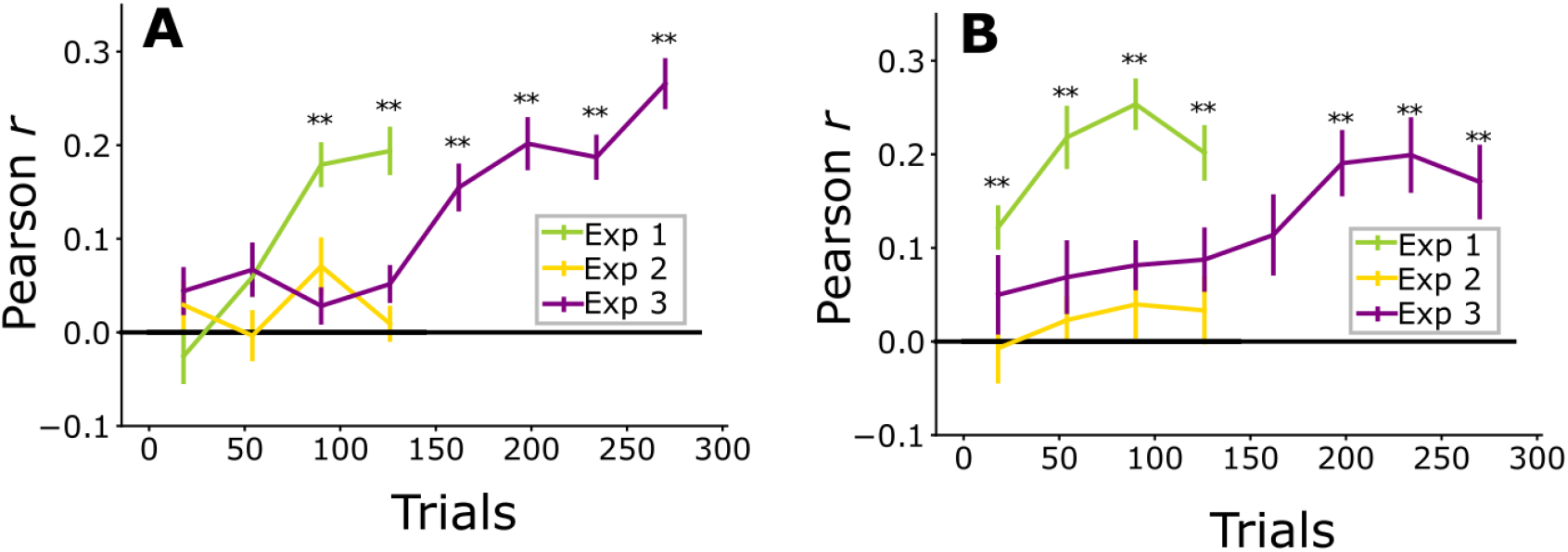
Changes in eye-movements due to acquired knowledge about the statistical structure of the stimulus have an increasingly direct link to performance in familiarity tests. Trial-by-trial eye-movement measures of each participant were correlated with individual learning success measured on the familiarity test. Single trial Pearson *r* values were averaged in successive 36-trial-long bins. **(A)** Exploratory gaze transitions successfully predicted performance on the familiarity test both in Exp. 1 and Exp. 3. Exploratory looking in all three experiments was not predictive of test performance in the initial bin, but it quickly emerged to a highly predictive level in Exp. 1, unlike in Exp. 2 and in the first half of Exp. 3, where Pearson *r* values remained at chance. However, in the second half of Exp. 3, a strong relationship between eye-movements and performance emerged matching that of Exp. 1. **(B)** Largely the same pattern of results was found with Confirmatory as with Exploratory transitions, with a faster emergence of statistical influence in Exp. 1. suggesting that returns could reflect a hypothesis testing process of learning. (Error Bars: SEM; ** *p*<.01 after Bonferroni correction).

**Figure 4.**
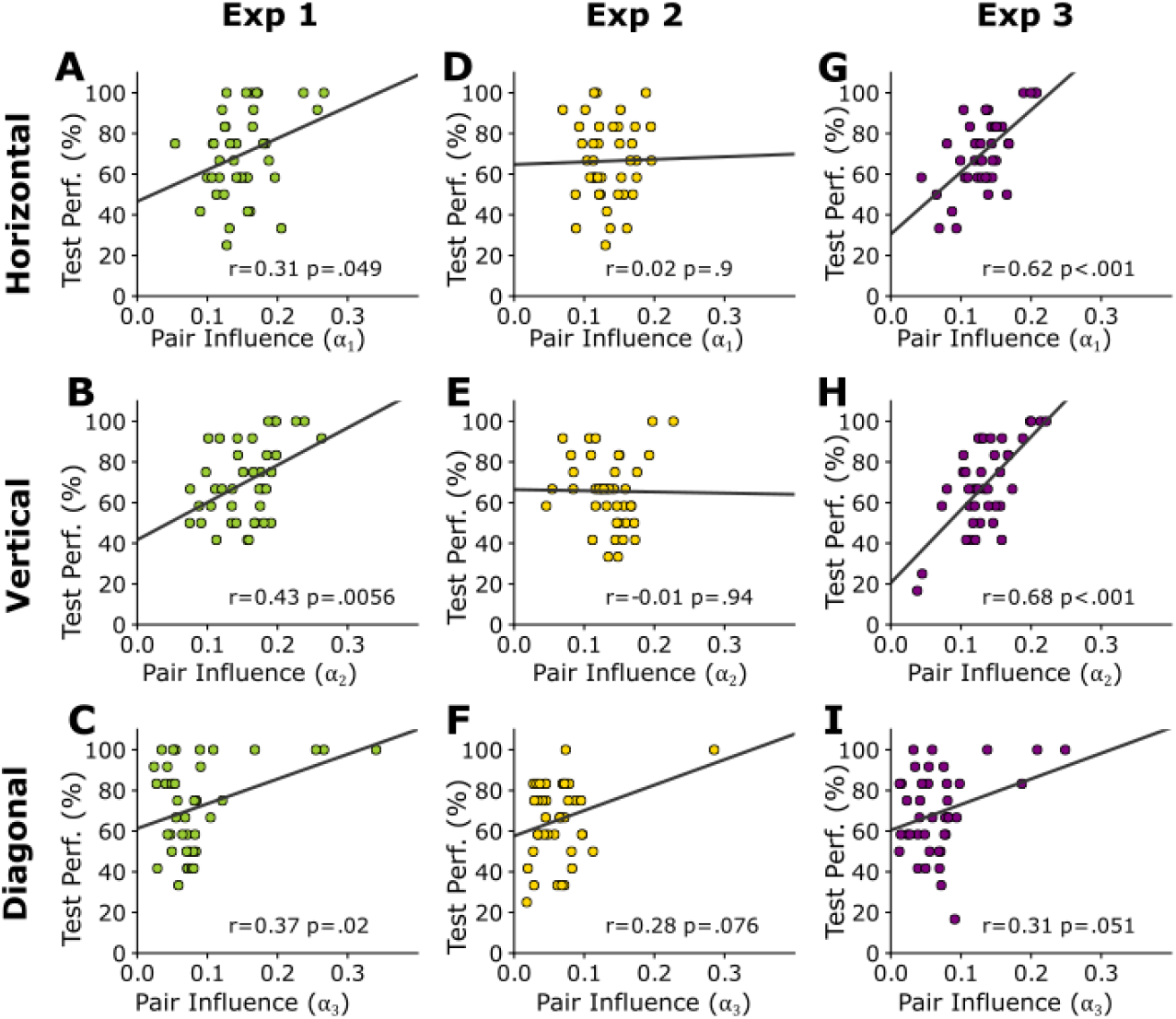
Familiarity test performance is predicted by eye-movement changes due to both implicit and explicit learning of stimulus regularities. On the x axes, parameters of the model-based analysis individually fitted to all gaze-transition data are shown, indicating how strongly a particular pair structure influenced eye-movements relative to the average exploration behavior of the participant. The model had three parameters, corresponding to Horizontal- (α_1_,**Top Row)**, Vertical- (α_2_, **Middle Row)**, Diagonal-pairs (α_3_ **Bottom Row)**, representing the relative increase in the number of looks that were in agreement with the spatial arrangement of the pairs. On the y axes, performance on the familiarity test trials containing true pairs from the corresponding orientation is presented. Pearson *r* and *p* and Least Square regression line are shown for each condition. The specific link between eye-movements and the content of learning was especially strong in Exp. 3 (Right Column), both for horizontal and vertical pairs. The same two directions also showed a significant relationship in Exp. 1 (Left Column), with a weaker relationship for diagonal pairs due to a stronger ceiling effect. None of the links were significant in Exp. 2 (Middle Column).

### Implicit learning of spatial regularities

In Experiment 1, we demonstrated a direct link between learning complex regularities and eye-movements when an explicit instruction provided a cognitive support for learning and visual explorations. In Experiments 2 and 3, we investigated whether this link between learning and eye-movements persists when people are solely exposed to the stimuli without any previous knowledge or instructions about regularities within the stimuli. Since learning could only be assessed without interference with implicitness after the end of the exposure period (by the familiarity test), we used two different training lengths in order to assess the link between the strength of learning and its influence on eye-movements at two different stages of learning. Participants demonstrated significant learning in the familiarity test in both experiments (Fig 1D; Exp. 2: t(39)= 6.81, *p*<.001, *d*=1.08; Exp. 3: t(39)= 7.58, *p*<.001, *d*=1.2), with the performance in Exp. 3 numerically above that in Exp. 2 (Exp. 3 69.65%, [64.64, 74.67] vs. Exp. 2 65.9% [61.38, 70.43]), but this difference was not statistically significant (*t*_78_=1.07, *p*= .286, d=.24, Bayes Factor= .38).

### Change in eye-movements during implicit learning

Analyzing the effect of the underlying structure on the eye-movements with least-square regression analysis, we found a striking contrast between the two experiments. In Exp. 2, we found no evidence conveyed by regression slopes of any increase in within-pair fixations rates either for exploratory (β=−0.0039, *p*=.513, Fig 2C) or for confirmatory looks (β=0.007, *p*=.643, Fig 2D). In contrast, and more similarly to Exp. 1, observers’ changing fixation rates in Exp. 3 reflected an increasing influence of the pair structure on eye movements over time both in exploratory (β=0.0068, *p*=.005, Fig 2E) and confirmatory looks (β=0.0139, *p*=.012, Fig 2F). Compensating the potential confounding effect of variable numbers of eye movements within trials, we reanalyzed the data with a Bayesian mixed model and confirmed the significance of the regression slope in Exp 3, and the lack of such effect in Exp. 2.

### Eye-movements predict implicit learning performance

In Exp. 2, the eye-movement measures were not predictive of the outcome of the familiarity test (Exploratory: r(38)=0.17, *p*=.308; Confirmatory *r*(38) = 0.18, *p=*.26). In contrast, in Exp. 3, both measures had a strong correlation with learning performance (Exploratory: r(38)=0.55, *p*<.001; Confirmatory *r*(38) =0.54, *p*<.001). This relationship between learning and eye-movements in Exp. 3 emerged gradually and revealed the strong link only by the second half of the experiment (Fig 3 A-B).

### Eye-movements specifically predict the content of learning

There was a similar difference between the two experiments in terms of the link between the orientation-specific changes of eye-movements (model α_1-3_) and familiarity test performance. Predictive relationships were absent in Exp. 2 (Fig 4 D-F), while in Exp. 3, there was a very strong relationship between the magnitude of orientation-specific influence on observer’s eye-movements and their pair-specific test performance. For both horizontal and vertical pairs, this effect was strong and highly significant (Fig 4 G-H), while for diagonal pairs, it was weaker and marginally significant (Fig 4 I). We confirmed that these correlations in Exp 3 were not due to general learning effects, but they were highly specific to the particular features the participants learned.

### Test similarity of learning influences

Although our results so far demonstrated that learning both explicitly and implicitly changed eye-movement patterns (Figure 2), it is unclear if these changes in eye-movement were linked only to the amount of learning regardless of whether this knowledge was acquired in an explicit or implicit manner. We hypothesized that, while eye-movements in Experiments 2 and the first half of Experiment 3 were obviously similar, in the second half of Experiment 3, when participants already gained some implicit knowledge of the structure of the input comparable to the gain from explicit instructions in Experiment 1, the eye-movement pattern changes would be indistinguishable for those in Experiment 1. To test this hypothesis, we performed four analyses of covariance (ANCOVA) comparing within-pair eye-movements between Exps 1 or 2 and the two halves of Exp 3, while controlling for the amount of learning. In these analyses, the *Average rate* of within-pair eye-movements of each participant combined across exploratory and confirmatory looks was the dependent variable. The *Type of the experiment* was the independent categorical variable, and *Test performance* indicating the amount of learning was the covariate with an interaction term between the covariate and the independent variable.

### The relationship of learning & eye-movements across experiments

Confirming the results in the sections above, the comparisons between Exps 1 and 3 showed that the “Test performance” covariate had a very strong influence on eye-movements (Exp1/Exp3-First half: F(1,76)=29.22, *p*<.001, *η*_*p*_^*2*^=.28; Exp1/Exp3-Second Half: F(1,76)=45.67, *p*<.001, *η*_*p*_^*2*^=.38). Comparing the first half of Exp 3 and Exp 1 (**Fig 5A**), we found that this influence of test performance had a significant interaction with the Type of experiment (F(1,76)=5.36, *p*=.023, *η*_*p*_^*2*^=.07), which rendered the lack of overall main effect of Type of experiment (F(1,76)=1.64, *p*=.204, *η*_*p*_^*2*^=.02) uninterpretable. By the second half of Exp. 3 (**Fig5C**), neither the slope F(1,76)=1.05, *p*=.31, *η*_*p*_^*2*^=.01), nor the overall eye-movements were dependent on the Type of the experiment (F(1,76)=2.04 *p*=.158, *η*_*p*_^*2*^=.03). Thus, gaze-patterns were strongly influenced by the learned knowledge and became indistinguishable between the explicit and implicit experimental conditions. The same analysis for Exp. 3 vs. Exp. 2 showed the opposite pattern. When comparing Exp 2 to the first half of Exp 3 (**Fig5B**), the Type of experiment had neither a significant main effect (F(1,76)=1.33, *p*=.253, *η*_*p*_^*2*^=.02) nor an interaction (F(1,76)=0.51, *p*=.477, *η*_*p*_^*2*^=.01) with the covariate Test performance confirming high similarity across the two conditions. In contrast, when comparing Exp 2 to the second half of Exp. 3 (**Fig5D**), although the Type of experiment had a significant main effect (F(1,76)= 10.67, *p*=.002, *η*_*p*_^*2*^=.12), it also had a significant interaction F(1,76)= 10.58, *p*=.002, *η*_*p*_^*2*^=.12) with Test performance indicating that very different causes shaped the gaze-patterns in the two experiments. Importantly, the influence of the Test performance covariate was significant already when the first half of Exp 3 was compared to Exp. 2 (F(1,76)=8.18, *p*=.005, *η*_*p*_^*2*^=.1), but as expected, it became stronger when the second half of Exp 3 was considered (F(1,76)=21.66, *p*<.001, *η*_*p*_^*2*^=.22). These analyses indicate that while initially the (relatively weak) relationship between eye-movements and the learning of the underlying structure was very similar between the first half of Exp. 3 and Exp 2, as implicit knowledge accumulated further in Exp 3, it started to influence eye-movements more strongly, and the eye-movement patterns in Exp 3 were influenced in the same way as in the completely explicit learning context of Exp. 1. Thus, these results confirm our hypothesis that in our experiments, the amount of the acquired knowledge is the main driving force behind the changes in eye-movements patterns regardless of the explicit or implicit nature of the experimental conditions.

**Figure 5.**
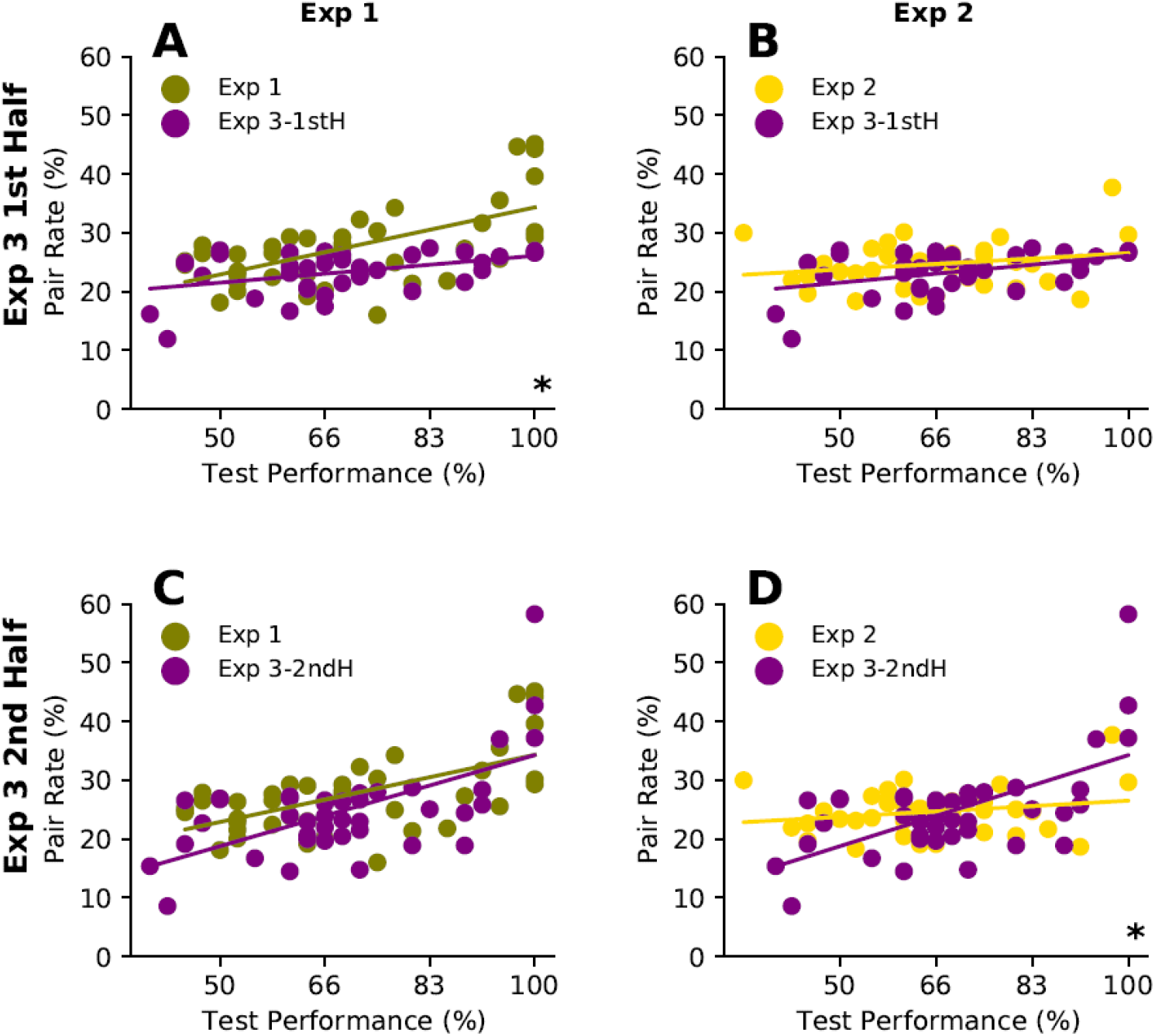
The relationship between learning and eye-movements during implicit and explicit learning becomes very similar over time. Scatter plots between familiarity test performance (x-axis) and the ratio of within-pair eye-movements (y-axis) across the three different experiments with Exp 3 splitted to two halves. *Top Row:* Comparison of the first half of Exp 3 to Exp 1 (A) and Exp 2 (B). *Bottom Row:* Comparison of the second half of Exp 3 to Exp 1 (C) and Exp 2 (D). Dots represent mean pair rate combined across Exploratory and Confirmatory looks and the corresponding test performance of individual participants. The lines indicate results of multiple linear regression corresponding to the ANCOVA described in the main text. Stars (**\*)** in the bottom right corners mark a significant interaction between test performance and the covariate Experiment type (A,D), the main focus of this analysis. The main effect of test-performance was significant in all four analyses.

### Statistical influence on eye-movements is automatic, but only emerges after sufficient learning

In Exp. 3, we found that, given sufficiently long exposure, learning regularities implicitly and learning them by explicit instructions (Exp. 1) influence visual exploration very similarly. Importantly, the relationship between acquired knowledge and gaze patterns was indistinguishable between the explicit and long implicit conditions. This suggests that the influence of learned knowledge of environmental statistics on eye-movements is automatic, and it does not require a well-defined task or cognitive awareness to emerge. We also found that this effect was tightly linked to the specific knowledge acquired about the statistics of the input. Meanwhile, in the shorter implicit experiment (Exp. 2), we found comparably large learning in the familiarity test without any detectable influence of this learning on eye movements. This provides evidence about the complex relationship between learning and eye movements indicating that precise assessment of the acquired knowledge and good sensitivity in measuring changes in eye-movement patterns will be needed for an ultimate characterization of their relation.

## Discussion

Using a novel gaze-contingent statistical learning paradigm, we clarified three aspects of how sensory learning and eye movement patterns interact. First, we confirmed that acquiring knowledge about the underlying structure of the visual environment accumulated by sensory learning can have an effect on the patterns of eye movements even on the short run. Second, we showed that this effect is highly specific to the statistical composition of the incoming sensory input as the knowledge acquired by learning and characterized by different orientations of the underlying chunks could be reliably identified by the individual looking patterns. Finally, we found that, apart from the learning speed, the effect of knowledge on eye movements was independent of whether the observers gained it using explicit instruction about the underlying structure of the input or they obtained it by exploring the scenes without any specific prior information.

Previous studies investigating the relationship between environmental regularities and eye-movements fall roughly into two groups investigating complementary aspects of the phenomenon. Studies in the first group focused on the interaction between explicitly or implicitly defined but already available internal knowledge and eye movements in various tasks by investigating the number and position of fixations necessary for finding a target in a display or making a decision(Chukoskie et al., 2013; Hoppe & Rothkopf, 2019; Morvan & Maloney, 2012; Najemnik & Geisler, 2005; Peterson & Kramer, 2001; Yang et al., 2017). Studies in the other group investigated the effect of learning on eye movements but only in terms of learning temporal regularities and adjusting the timing of fixations accordingly(Glimcher, 2003; Hoppe & Rothkopf, 2016). Our study is the first to combine these two aspects by investigating the ongoing process of developing an internal representation of the input’s spatial structure and showing how the momentary result of this learning process continuously interacts with the pattern of eye movements.

Our design also allowed addressing directly the controversial issue regarding the role of explicit vs. implicit knowledge in controlling eye movements. Some studies found that only explicit memories have an influence on eye-movements(Hannula et al., 2012; Smith et al., 2006), while others reported that eye-movements can be used to detect memory traces that are not yet amenable to conscious report(Hannula et al., 2012; Hannula & Ranganath, 2009). Although our findings do not decisively resolve this controversy, we show in a unified setup how eye-movements can reflect memory traces as an outcome of both an explicit and a sufficiently long implicit learning process. While there is an ongoing semantic debate about the definition of explicit vs. implicit memory(Batterink et al., 2015; Greenwald & Banaji, 2017; Roediger, 1990), our results provide two important observations pertinent to the issue. First, the majority of the implicit observers even in the long implicit experiment (Exp 3) did not have explicit access to the gained knowledge about the underlying scene, as indicated by their verbal post-test report. Nevertheless, they showed a monotonic increase in correlation between their implicitly-acquired knowledge and effects on their eye movements. Moreover, even after removing subjects, who performed perfectly on the familiarity test in Exp 3, in a control measure, our conclusion remained the same. Second, the nature of changes in the eye movements in the implicit and explicit conditions were very similar as measured by the rate of exploratory and confirmatory gaze switches, suggesting that the underlying processes were also shared across the two conditions. These two observations indicate that the phenomenon we uncovered is, indeed, a general and automatic process that is driven by knowledge regardless of whether this knowledge is acquired implicitly or explicitly. Moreover, this automatic process influences information collection during perception continuously and in proportion to the amount of acquired knowledge.

While the correlational nature of our findings does not allow the assessment of the causal relationship between eye-movements and learning, our results have implications for the suitable framework for capturing the interplay between learning new information and selective data acquisition due to eye movements constrained by internal knowledge. The continuous ongoing nature of the emerging knowledge-based effect on eye movements and the independence of this knowledge of explicitness does not support frameworks positing that eye-movements are affected in an attention-like manner only when the underlying structure of the environment has been learned and it is explicitly accessible. Based on our results, it is more parsimonious to assume an ongoing bi-directional relationship, in which learning influences eye-movements and eye-movements scaffold learning, reflecting a continuous intertwined link between new sensory input and top-down memory related control(Chun & Turk-Browne, 2007; Gottlieb, 2012). Computationally, this process is better represented by dynamically evolving hierarchical inference making(Lake et al., 2015), in which prior knowledge and momentarily collected information is jointly handled for continuously interpreting and controlling sensory input than by two-stage schemes with an initial sweep of bottom-up process followed by specific top-down cognitive filtering(Itti & Baldi, 2009; Schütz et al., 2012).

Finally, the method we used in this study can also improve our understanding of the computations involved in statistical learning. Despite being considered as a fundamental form of human knowledge gathering, statistical learning is still not well understood at the process level. This is due to the fact that the majority of studies use the early methodology established in classical papers (Fiser & Aslin, 2001; Saffran et al., 1999), in which the effect of learning is measured on a separate test phase following exposure and this provides only limited information about characteristics of learning (Siegelman et al., 2017). Although recently, different methods were proposed to deal with this problem by tracking the ongoing processes of learning visual regularities (Karuza et al., 2014; Siegelman et al., 2017, 2018), these methods are restricted to temporal statistical learning and raise new concerns due to using explicit instructions instead of truly implicit learning, and increased number of test trials that could interfere with learning. In contrast, our method can be used to track the learning of complex spatial regularities in a natural manner as in the classical experiments, since it relies on an independent and unconscious behavioral measure -eye movement patterns- that does not require changing the original setup of statistical learning and still provides information continuously about the characteristics of the emerging representation.

In conclusion, we provided evidence for the first time for a continuous and tight link between human visual information sampling strategies manifested by eye movements and the emerging internal knowledge of environmental regularities. Our results frame natural vision as a process, in which active selection from the incoming information and internal knowledge jointly determine both the interpretation of the input and further changes in internal knowledge.

## Methods

### Participants

Altogether 120 participants naïve about the purpose of the study and about statistical learning were recruited via a local student organization and received monetary compensation for their participation. 40 participants were assigned to each of the three experiments (Exp. 1: age: 25.5 +/− 4.6 years, 13 male; Exp. 2: age: 22.1 +/− 2.8 years, 13 male; Exp. 3: age: 23 +/− 5.5 years, 10 male). We chose a sample size larger than most previous statistical learning studies(Batterink et al., 2015; Fiser & Aslin, 2001; Turk-Browne et al., 2005) based on power analysis, as we wanted to assess the variability in the individual learning performances. One additional participant completed Exp. 2 but was excluded from the final sample, because upon completing the study revealed not being naïve about visual statistical learning.

### Procedure

In Experiment 1, after calibration and practice, but before the start of the main experiment, participants were instructed to explore the scenes and find pairs of shapes that always appear next to each other in a horizontal, vertical or diagonal arrangement. They were also told that they would be questioned about the identity of the pairs afterwards (Explicit instructions). Participants had 6 seconds to explore each of the 144 scenes, presented in a random order, resulting in a total training time of approximately 16 minutes.

All aspects of Experiments 2 and 3 were identical to those in Experiment 1 except for the lack of explicit instructions. After calibration and practice, but before the start of the main experiment, participants were told to explore the scenes and pay attention to what they see. They were also told that they will be tested on what they had seen after the exploration phase, however, they were not told about any potential regularity or structure in the stimuli nor about the nature of the subsequent test. These are the canonical conditions of implicit visual statistical learning used in previous studies(Fiser & Aslin, 2001; Turk-Browne et al., 2005). Exp. 2 was the same length as Exp. 1 (~16 mins), but in Exp. 3, the learning phase was double in length: each one of the 144 unique scenes were presented once in each half of the experiment in a different random order. In Exp. 3, completing the learning phase took approximately 32 mins, with a short break in the middle, where participants were kindly asked to continue paying attention.

All experiments were conducted in a dimly lit and sound attenuated room. A Tobii EyeX 60Hz eye-tracker was calibrated using a seven-point calibration from a viewing distance of 60 cm. After calibration, participants completed ten 6-second-long practice trials, where randomly selected images of dogs were revealed in a gaze-contingent manner within the 3 × 3 grid: the content of each cell was visible only when the observer’s gaze fell within the central 5.7 × 5.7 degrees of the cell in two subsequent eye position samples (taken approx. 15 ms apart), otherwise the given cell was shown empty. The trials in the learning phase of each experiment were also 6-second-long and they followed the same gaze contingent rule as during practice.

Each trial started by a fixation cross appearing in one of the empty grid cells, where the observer had to fixate to initiate the trial. The position of the fixation cross was uniformly distributed across trials, appearing at the center of each cell of the 3 × 3 grid an equal number of times during the experiment in a random order. Unlike previous spatial statistical learning studies, the full scenes in these trials were never visible at once. Instead, individual shapes were revealed in a gaze-contingent manner, when the participants’ gaze was inside the mid-region of a cell. When participants looked at a cell containing the shape, the shape appeared at full contrast as long as the participant’s gaze was in the given cell, but gradually faded away becoming invisible within 1.5 sec when the participant looked away to a different cell. This way, maximally two shapes of the scene were displayed at any given time and only one of them at full contrast. If the observer’s gaze was in the mid-region of a cell not containing a shape in a given trial, a gray rectangle was revealed indicating that the cell was empty in order to reduce the observer’s uncertainty whether s/he managed to fixate on the cell. These gray rectangles remained visible until the trial was over, thereby ensuring that the end of each trial was easily noticeable. Participants were free to visit or revisit with their gaze any of the cells during the trial. When the trial was over after 6 seconds, all shapes and gray rectangles disappeared, and after a 500ms inter-trial-interval, the next fixation-cross appeared at one of the cells to initiate the start of the next trial.

At the end of the learning phase, after a short break, a two-interval-forced-choice (2-IFC) test session followed, with trials in which participants were told to select the more familiar of the two pair combinations presented based on what they had seen during the learning phase. For the test, 6 foil pairs (with two shapes that never appear in the presented arrangement during learning) were created from the original shapes and those were tested in a fully counterbalanced manner against each of the real pairs of the inventory, resulting in 36 test trials presented in a random order. The within-test trial order of the real versus foil pair was pseudo-randomly balanced across the test. On each trial, participants used the left and right arrow keys for the 1^st^ and 2^nd^ pair, respectively, to indicate which pair was more familiar.

### Data Analysis & Measures

All data were analyzed in Python, and statistics were calculated using the SciPy, *scikit*-learn, *Pingouin* and *statsmodels* libraries. Bayes factors were calculated using the method proposed by Rouder et al (Rouder et al., 2009) with an uninformative prior. Since the exact gaze position within the central region of each cell had no functional consequence, eye-movement data was analyzed based on whether or not the fixation samples were within the gaze contingent central region of one of the cells. On average, participants made more than seven (7.2 +/− 1) transitions between the central regions of different cells in a trial. From these transition events, we focused on the ones that were potentially related to learning by using a method detailed below. Since the number of transitions could also change as the learning session progressed, we focused on proportions and not on the absolute number of events.

Eye-movement transition data was separated into two different measures that could indicate different behaviors: *exploratory transitions* and *confirmatory returns*. An exploratory transition was defined as a gaze transition to a cell for the first time during a trial, while a confirmatory return was defined as transition to a cell that had already been visited on the current trial. The difference between these events is important, since in case of a return, the participant could be more certain what s/he would see at a given location, as s/he had already seen the content within the last few seconds. In case of an exploratory transition, no such information was available, therefore, the content of the cell could be predicted/expected only if 1) the cell contained a member of a shape pair whose other member the participant already saw during the current trial, AND 2) only if the participant had already learned about the spatial relationships between shapes during the previous trials. Within the exploratory and confirmatory measures, we calculated the proportion of looks that were performed from a shape to its pair, and used this calculation for the assessment of whether the underlying statistical structure had an effect on the transitions. Finally, as a combined measure, we used the rate of within pair eye-movements, which was defined as the proportion of gaze transitions when participants looked from a shape to the cell containing its pair as opposed to other cells.

For the analysis of temporal changes in the gaze data across trials, we used regression to predict the eye-movement data with trial number as a predictor. We analyzed the results with two different regression methods, and found support with both of them for the same conclusions. The first method was a simple linear regression predicting the average eye-movement measures across participants (Fig 2). The second was a linear mixed model, predicting a slope for eye-movements across participants, but including a random intercept for each observer.

To analyze whether temporal changes in looking behavior across trials were linked to learning (Fig 3), we calculated the Pearson correlation between our eye-movement measures on each trial and the performance in the final familiarity test. Next, we divided the obtained *r* values in 36 trial-long consecutive bins (yielding 4 bins in Exps. 1 & 2 and 8 bins in Exp. 3), and analyzed whether the *r* values in each bin were different from zero using a standard one sample t-test. For statistical correction of multiple comparisons, the Bonferroni method was used.

### Computational Analysis

Our goal was to quantify how much participants’ gaze trajectories changed from random exploration to a pattern determined by statistical regularities over the duration of the experiment. We used a model-based analysis to obtain a measure that could be fitted to all gaze transitions without relying on the selection of particular events. For each participant, the model measured the increase of alignment between looking behavior and the statistical structure of the stimuli compared to the average behavior as quantified by the distribution of transition probability across the cells of the grid. Since there were three types of regularities in the stimuli (link across horizontal, vertical, and diagonal orientations), the model had three parameters (α_1-3_), representing increased gaze transitions between shapes forming pairs in each of the three orientations. For example, the value of α_1_ represented an increased probability of looking from shape1, which was a member of a horizontal pair, to the position of shape2, the other shape in the pair. For each observer, the values of the three parameters were fitted trial-by-trial using the maximum likelihood method. To test whether these orientation-specific changes in eye movement behavior during the learning phase could predict performance in the test session, we separated the 36 test trials based on the orientation of the true pair in the trial, yielding 12 test trials for each orientation. Next, we used Pearson correlation to predict orientation specific test performance based on the fitted model parameters of each participant (Fig 4).

## References

Batterink, L. J., Reber, P. J., Neville, H. J., & Paller, K. A. (2015). Implicit and explicit contributions to statistical learning. Journal of Memory and Language, 83, 62–78.

Brockmole, J. R., & Henderson, J. M. (2006). Using real-world scenes as contextual cues for search. In Visual Cognition (Vol. 13, Issue 1, pp. 99–108). https://doi.org/10.1080/13506280500165188

Chukoskie, L., Snider, J., Mozer, M. C., Krauzlis, R. J., & Sejnowski, T. J. (2013). Learning where to look for a hidden target. Proceedings of the National Academy of Sciences of the United States of America, 110 Suppl 2, 10438–10445.

Chun, M. M., & Turk-Browne, N. B. (2007). Interactions between attention and memory. Current Opinion in Neurobiology, 17(2), 177–184.

Findlay, J. M., & Gilchrist, I. D. (2003). Active Vision: The Psychology of Looking and Seeing. Oxford University Press.

Fiser, J., & Aslin, R. N. (2001). Unsupervised statistical learning of higher-order spatial structures from visual scenes. Psychological Science, 12(6), 499–504.

Glimcher, P. W. (2003). The neurobiology of visual-saccadic decision making. In Annual Review of Neuroscience (Vol. 26, Issue 1, pp. 133–179). https://doi.org/10.1146/annurev.neuro.26.010302.081134

Gottlieb, J. (2012). Attention, learning, and the value of information. Neuron, 76(2), 281–295.

Greenwald, A. G., & Banaji, M. R. (2017). The implicit revolution: Reconceiving the relation between conscious and unconscious. The American Psychologist, 72(9), 861–871.

Hannula, D. E., Baym, C. L., Warren, D. E., & Cohen, N. J. (2012). The eyes know: eye movements as a veridical index of memory. Psychological Science, 23(3), 278–287.

Hannula, D. E., & Ranganath, C. (2009). The eyes have it: hippocampal activity predicts expression of memory in eye movements. Neuron, 63(5), 592–599.

Hayhoe, M., & Ballard, D. (2005). Eye movements in natural behavior. Trends in Cognitive Sciences, 9(4), 188–194.

Henderson, J. M., Hayes, T. R., Rehrig, G., & Ferreira, F. (2018). Meaning Guides Attention during Real-World Scene Description. Scientific Reports, 8(1), 13504.

Hoppe, D., & Rothkopf, C. A. (2016). Learning rational temporal eye movement strategies. Proceedings of the National Academy of Sciences of the United States of America, 113(29), 8332–8337.

Hoppe, D., & Rothkopf, C. A. (2019). Multi-step planning of eye movements in visual search. Scientific Reports, 9(1), 144.

Itti, L., & Baldi, P. (2009). Bayesian surprise attracts human attention. Vision Research, 49(10), 1295–1306.

Karuza, E. A., Farmer, T. A., Fine, A. B., Smith, F. X., & Jaeger, T. F. (2014). On-line measures of prediction in a self-paced statistical learning task. Proceedings of the Annual Meeting of the Cognitive Science Society, 36(36).

Kowler, E. (2011). Eye movements: the past 25 years. Vision Research, 51(13), 1457–1483.

Lake, B. M., Salakhutdinov, R., & Tenenbaum, J. B. (2015). Human-level concept learning through probabilistic program induction. Science, 350(6266), 1332–1338.

Land, M. F., & McLeod, P. (2000). From eye movements to actions: how batsmen hit the ball. Nature Neuroscience, 3(12), 1340–1345.

Li, C.-L., Pilar Aivar, M., Tong, M. H., & Hayhoe, M. M. (2018). Memory shapes visual search strategies in large-scale environments. In Scientific Reports (Vol. 8, Issue 1). https://doi.org/10.1038/s41598-018-22731-w

Mack, S. C., & Eckstein, M. P. (2011). Object co-occurrence serves as a contextual cue to guide and facilitate visual search in a natural viewing environment. Journal of Vision, 11(9), 1–16.

Morvan, C., & Maloney, L. T. (2012). Human visual search does not maximize the post-saccadic probability of identifying targets. PLoS Computational Biology, 8(2), e1002342.

Najemnik, J., & Geisler, W. S. (2005). Optimal eye movement strategies in visual search. Nature, 434(7031), 387–391.

Nelson, J. D., & Cottrell, G. W. (2007). A probabilistic model of eye movements in concept formation. Neurocomputing, 70(13-15), 2256–2272.

Peterson, M. S., & Kramer, A. F. (2001). Attentional guidance of the eyes by contextual information and abrupt onsets. Perception & Psychophysics, 63(7), 1239–1249.

Roediger, H. L., 3rd. (1990). Implicit memory. Retention without remembering. The American Psychologist, 45(9), 1043–1056.

Rouder, J. N., Speckman, P. L., Sun, D., Morey, R. D., & Iverson, G. (2009). Bayesian t tests for accepting and rejecting the null hypothesis. Psychonomic Bulletin & Review, 16(2), 225–237.

Saffran, J. R., Johnson, E. K., Aslin, R. N., & Newport, E. L. (1999). Statistical learning of tone sequences by human infants and adults. Cognition, 70(1), 27–52.

Schütz, A. C., Trommershäuser, J., & Gegenfurtner, K. R. (2012). Dynamic integration of information about salience and value for saccadic eye movements. Proceedings of the National Academy of Sciences of the United States of America, 109(19), 7547–7552.

Siegelman, N., Bogaerts, L., & Frost, R. (2017). Measuring individual differences in statistical learning: Current pitfalls and possible solutions. Behavior Research Methods, 49(2), 418–432.

Siegelman, N., Bogaerts, L., Kronenfeld, O., & Frost, R. (2018). Redefining “Learning” in Statistical Learning: What Does an Online Measure Reveal About the Assimilation of Visual Regularities? Cognitive Science, 42 Suppl 3, 692–727.

Smith, C. N., Hopkins, R. O., & Squire, L. R. (2006). Experience-dependent eye movements, awareness, and hippocampus-dependent memory. The Journal of Neuroscience: The Official Journal of the Society for Neuroscience, 26(44), 11304–11312.

Spalek, T. M., & Hammad, S. (2005). The left-to-right bias in inhibition of return is due to the direction of reading. Psychological Science, 16(1), 15–18.

Turk-Browne, N. B., Jungé, J., & Scholl, B. J. (2005). The automaticity of visual statistical learning. Journal of Experimental Psychology. General, 134(4), 552–564.

Võ, M. L.-H., & Wolfe, J. M. (2013). The interplay of episodic and semantic memory in guiding repeated search in scenes. Cognition, 126(2), 198–212.

Wolfe, J. M., & Horowitz, T. S. (2017). Five factors that guide attention in visual search. In Nature Human Behaviour (Vol. 1, Issue 3). https://doi.org/10.1038/s41562-017-0058

Yang, S. C.-H., Lengyel, M., & Wolpert, D. M. (2017). Correction: Active sensing in the categorization of visual patterns. eLife, 6. https://doi.org/10.7554/eLife.25660

Yarbus, A. L. (1967). Eye Movements During Perception of Moving Objects. In Eye Movements and Vision (pp. 159–170). https://doi.org/10.1007/978-1-4899-5379-7_7

